# Cortical networks for encoding near and far space in the non-human primate

**DOI:** 10.1101/261677

**Authors:** Justine Cléry, Olivier Guipponi, Soline Odouard, Claire Wardak, Suliann Ben Hamed

**Affiliations:** Institut des Sciences Cognitives Marc Jeannerod, UMR5229, CNRS-Université Claude Bernard Lyon I, 67 Boulevard Pinel, 69675 Bron, France

**Keywords:** near space, peripersonal space, far space, macaque monkey, fMRI

## Abstract

While extra-personal space is often erroneously considered as a unique entity, early neuropsychological studies report a dissociation between near and far space processing both in humans and in monkeys. Here, we use functional MRI in a naturalistic 3D environment to describe the non-human primate near and far space cortical networks. We describe the co-occurrence of two extended functional networks respectively dedicated to near and far space processing. Specifically, far space processing involves occipital, temporal, parietal, posterior cingulate as well as orbitofrontal regions not activated by near space, possibly subserving the processing of the shape and identity of objects. In contrast, near space processing involves temporal, parietal and prefrontal regions not activated by far space, possibly subserving the preparation of an arm/hand mediated action in this proximal space. Interestingly, this network also involves somatosensory regions, suggesting a cross-modal anticipation of touch by a nearby object. Last, we also describe cortical regions that process both far and near space with a preference for one or the other. This suggests a continuous encoding of relative distance to the body, in the form of a far-to-near gradient. We propose that these cortical gradients in space representation subserve the physically delineable peripersonal spaces described in numerous psychology and psychophysics studies.

**Highlights:**

- Near space processing involves temporal, parietal and prefrontal regions.
- Far space activates occipital, temporal, parietal, cingulate & orbitofrontal areas.
- Most regions process both far & near space, with a preference for one or the other.
- Far-to-near gradient may subserve behavioral changes in peripersonal space size.

## INTRODUCTION

Our environment is often perceived as a unitary space, however growing evidence demonstrates that the brain contains a modular and a dynamic representation of space. Early neuropsychological reports demonstrate that the unilateral ablation of the frontal eye fields produces, in monkeys, an inattention to contralateral objects, more pronounced for far than for near objects (Rizzolatti et al., 1983). In contrast, the inattention to contralateral objects produced by the unilateral ablation of premotor area 6 is more pronounced for near than for far objects (Rizzolatti et al., 1983). In 1991, a single case study presents the first neuropsychological evidence for a left neglect in near space but not in far space after a unilateral right hemisphere stroke in humans (Halligan and Marshall, 1991). The finding of the opposite dissociation confirms that, as for monkeys, far and near space are separately coded by the human brain (Cowey et al., 1994, 1999; Vuilleumier et al., 1998), though a task dependence of far and near space processing deficits is reported (Aimola et al., 2012; Keller et al., 2005).

In recent years, fMRI studies show that the coding of near space involves a dorsal network including the left dorsal occipital and intraparietal cortex and the left ventral premotor cortex, while the coding of far space involves a ventral network including the ventral occipital cortex bilaterally and the right medial temporal cortex (Aimola et al., 2012; Weiss et al., 2000).

Peripersonal neurons fire both when a tactile stimulus is delivered to the animal’s skin and when a visual stimulus is presented in the space near the part of the body where the tactile field is located. These have been described in several cortical regions: the prefrontal cortex (Fogassi et al., 1996; Gentilucci et al., 1988; Graziano et al., 1994; Gross and Graziano, 1995); in the parietal cortex where Hyvärinen and Poranen (1974) describe the visual response of parietal neurons “as an anticipatory activation” that appears before the neuron′s tactile RF is touched. The multimodal ventral intraparietal area VIP stands out in this respect (Duhamel et al., 1998; Avillac et al., 2004, 2007, Guipponi et al., 2013, 2015a; Wardak et al., 2016). This region encodes both large field visual movement mimicking the consequences of the displacement of a subject within its environment (Bremmer et al., 2000, 2002a, 2002b) and the movement of visual objects within the near peripersonal space (Bremmer et al., 1997, 2013). A first study did not identify a sensitivity to the 3-dimensional structure of static stimuli in VIP (Durand et al., 2007), however a more recent study shows that VIP is involved in depth-structure processing (Van Dromme et al., 2016). In comparison, human VIP shows no preference for any particular spatial range (Quinlan and Culham, 2007).

Beyond these two prefrontal and parietal cortical regions, little is known about the whole brain network that is involved in far and near space processing in monkeys. In particular, while several functional magnetic resonance imaging (fMRI) studies describe the cortical regions involved in the processing of 3-dimensional shape from disparity (Durand et al., 2009; Joly et al., 2009) or shading and texture (Nelissen et al., 2009), to our knowledge, no study to date provides, in this specie, a description of the cortical networks involved in the processing of far and near space. fMRI performed in monkeys bridges the gap between human fMRI and monkey invasive methodologies, and allows to identify and localize the activations in specific voxels, beyond cytoarchitectonic definitions of functional areas to guide future electrophysiological recordings. Here, we use fMRI in monkeys immersed in a naturalistic 3D environment to describe a functional gradient going from selective near space coding, to preferential near space coding, to unselective space coding, to preferential far space coding and selective far space coding. We also describe a cortical network activated by intrusion into peripersonal space. Interestingly, this network is only partially overlapping with the near space coding network. Overall, these observations argue for multiple space representations the functional significance of which remains to be assigned.

## MATERIAL AND METHODS

All procedures were in compliance with the guidelines of European Community on animal care (European Community Council, Directive No. 86–609, November 24, 1986). All the protocols used in this experiment were approved by the animal care committee (Department of Veterinary Services, Health & Protection of Animals, permit number 69 029 0401) and the Biology Department of the University Claude Bernard Lyon 1. The animals’ welfare and the steps taken to ameliorate suffering were in accordance with the recommendations of the Weatherall report, "The use of non-human primates in research".

### Subjects and experimental setup

Two rhesus monkeys (female MZ, male MT, 10-8 years old, 6-10 kg) participated to the study. The animals were implanted with a plastic MRI compatible headset covered by dental acrylic. The anesthesia during surgery was induced by Zoletil (Tiletamine-Zolazepam, Virbac, 15 mg/kg) and followed by Isoflurane (Belamont, 1-2%). Post-surgery analgesia was ensured thanks to Temgesic (buprenorphine, 0.3 mg/ml, 0.01 mg/kg). During recovery, proper analgesic and antibiotic coverage were provided. The surgical procedures conformed to European and National Health guidelines for the care and use of laboratory animals.

During the scanning sessions, monkeys sat in a sphinx position in a plastic monkey chair positioned within a horizontal magnet (1.5-T MR scanner Sonata; Siemens, Erlangen, Germany). Their head was restrained and they were equipped with MRI-compatible headphones customized for monkeys (MR Confon GmbH, Magdeburg, Germany). A radial receive-only surface coil (10-cm diameter) was positioned above the head. Monkeys were required to fixate a LED placed at 90 cm away from their face, at eye level, aligned with their sagittal axis. Eye position was monitored at 120 Hz during scanning using a pupil-corneal reflection tracking system (Iscan^®^, Cambridge, MA). The calibration procedure involved the fixation LED and 4 additional LEDs, placed in the same coronal plane as the fixation LED. All five LEDs were sequentially switched on and off and the monkey was rewarded for orienting and maintaining its gaze towards the illuminated LED. These four additional LEDs were subsequently removed during the main task during which only the central LED was present. Monkeys were rewarded with liquid dispensed by a computer-controlled reward delivery system (Crist^®^) thanks to a plastic tube coming to their mouth. The reward probability and quantity increased as fixation duration increased according to a subject-specific schedule, thus positively reinforcing fixation behavior. Fixation was considered as successful when the eyes remained in a window of 1.5° around the fixation LED. The reward schedule was uncorrelated with the scanning schedule. The task and all the behavioral parameters were controlled by two computers running Matlab^^®^^ and Presentation^®^. Monkeys were trained in a mock scan environment approaching to the best the actual MRI scanner setup. Actual scanning was performed once their fixation performance was maximized (>85-90% of time inside the tolerance window).

### Task and stimuli

The animals were trained to maintain fixation on the central LED. This allowed to control for eye vergence signals all throughout the experimental runs. An enriched stable visual scene was installed around the fMRI aperture, in front of the monkeys, so as to maximize depth cues (2m high lateral black curtains hiding the experimenters, large wooden static sticks placed along the curtains and in the back of the room, visible to the monkeys throughout the experiment). Once this behavior was stabilized, monkeys were further trained to maintain fixation while 3D objects were presented at either an average of 15 cm in front of their eyes (i.e. in the space between their head and the fixation LED, see Figure 1C) or at an average of 150 cm (i.e. in the space beyond the fixation LED, see Figure 1C), thus allowing to stimulate the space situated respectively near and far from the animals. Stimulations were achieved with either a small cube (3x3x3 cm) or a large cube (30x30x30 cm, Figure 1A), attached to a rigid holding stick. These cubes had the same apparent size when the small cube was placed at 15cm from the subject and the larger cube was placed at 150cm (Figure 1C), thus allowing to control for size effects. These cubes were constructed with transparent plastic material and the presentation of the near small object did not hide the fixation LED from the monkey. In order to maximize depth cues, the edges of the cubes were highlighted with red stripes and their transparent faces were ornamented with fractal pictures, resized for each cube such that the edges and the images occupied the same proportion in each cube (Figure 1A). Each cube was attached to a thin wooden stick by means of a transparent nylon cord. During the stimulation duration, the cube was continuously agitated by the experimenter so as to prevent neuronal habituation, but only the cube and the stick were visible to the monkeys, as the experimenters stayed hidden behind the curtains. To confine visual stimulation to a depth range, marks were placed on the floor, on the sticks holding the cubes as well as on the curtains hiding the experimenter. Experimenter manipulating the large cube controlled the manipulation of the small cube by the other experimenter and vice versa. Experimenters flipped from large cube to small cube manipulation from one scanning session to the other, so as to minimize experimental systematic biases. In this context, and given the size of the cubes and the fact that these were agitated in all directions, the localization error of the cubes can be considered as marginal. Three conditions were tested in blocks of 13 pulses: 1) the small cube presented in the near space; 2) the small cube presented in the far space and 3) the big cube presented in the far space. Both during training and testing, the cubes were approached (1 pulse), agitated (13 pulses) and withdrawn (1 pulse) from the target location (near or far space) by two experimenters out of the field of view (behind the black opaque curtains), one controlling the small cube and the other the large cube. The stimulation instructions (count down, type of stimulation, pulse counts etc.) were delivered to them on a computer screen placed next to them and coupled to the experimental control system.

**Figure 1:**
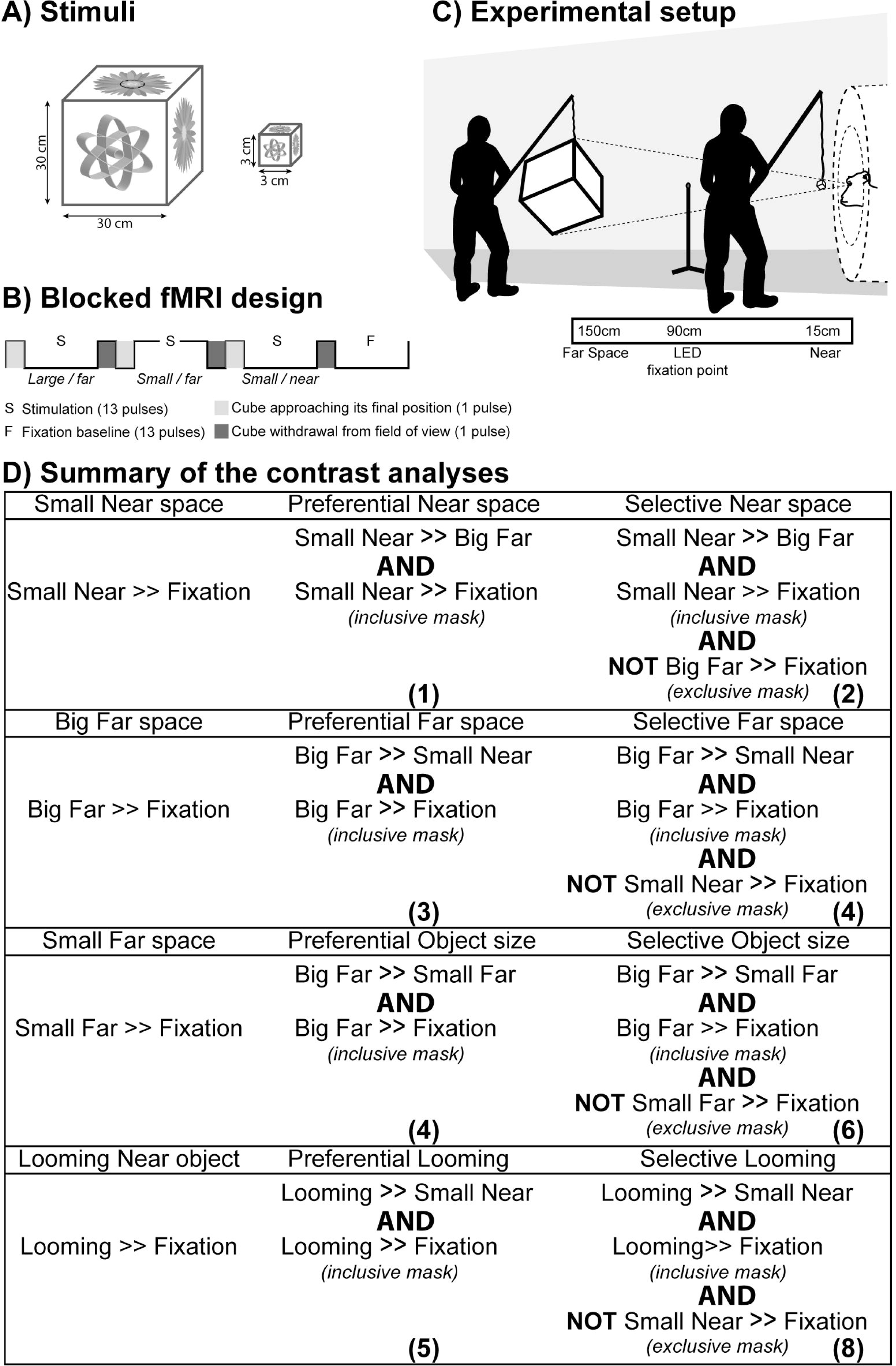
Experimental fMRI protocol. A) 3D naturalistic stimuli, a large 30x30x30 cm3 cube and an identical small 3x3x3 cm3 cube. The 6 faces of each transparent cube were decorated with 6 different fractal images. B) Block design. C) Experimental set up. Near space (15 cm from the monkey’s face) and far space (150 cm from the monkey’s face) were stimulated with the two types of 3D stimuli (same apparent sizes). Fixation is achieved at an intermediate position (red fixation LED, at 90cm). D) Summary of contrast analyses performed to extract the networks of interest: condition of interest higher than fixation (first column), preferential coding of condition of interest (second column) and selective coding of condition of interest (third column).

Functional time series (runs) were organized as follows (Figure 1B): a 13-volume block of stimulation category 1 was followed by a 13-volume block of stimulation category 2, a 13-volume block of stimulation category 3, and a 13-volume block of pure fixation (baseline). Before the beginning (resp. after the end) of each block of stimulation, 1 pulse was dedicated to the approach (resp. withdrawal) of the appropriate cube towards (resp. away from) the target space. Approaching or withdrawing the cubes involved only a minimal displacement of the curtains, thanks to two vertical slits in the curtains, placed respectively close to the magnet bore or at 150cm from the monkey. These slits were firmly closed back during the main stimulation blocks. A given sequence was played three times, resulting in a 174-volume run. The blocks for the 3 categories were presented in 6 counterbalanced possible orders.

A previous study involving both these monkeys (Guipponi et al., 2015a) and using a classical retinotopy localizer (Fize et al., 2003), allowed us to generate individual whole brain maps of center vs. periphery visual coding (see Supplementary data S1).

### Scanning

Before each scanning session, a contrast agent, monocrystalline iron oxide nanoparticle (Feraheme^®^, Vanduffel et al., 2001), was injected into the animal’s femoral/saphenous vein (4-10 mg/kg). Brain activations produce increased BOLD signal changes. In contrast, when using MION contrast agents, brain activations produce decreased signal changes (Kolster et al., 2014). For the sake of clarity, the polarity of the contrast agent MR signal changes, which corresponds essentially to a cerebral blood volume (CBV) measurement, was inverted. We acquired gradient-echo echoplanar (EPI) images covering the whole brain (1.5 T; repetition time (TR) 2.08 s; echo time (TE) 27 ms; 32 sagittal slices; 2x2x2-mm voxels). A total of 34 (22) runs was acquired for MZ (/MT).

### Analysis

A total of 20 (15) runs were selected based on the quality of the monkeys’ fixation throughout each run (>80% within the tolerance window). Time series were analyzed using SPM8 (Wellcome Department of Cognitive Neurology, London, United Kingdom). For spatial preprocessing, functional volumes were first realigned and rigidly coregistered with the anatomy of each individual monkey (T1-weighted MPRAGE 3D 0.6x0.6x0.6 mm or 0.5x0.5x0.5 mm voxel acquired at 1.5T) in stereotactic space. The JIP program (Mandeville et al., 2011) was used to perform a non-rigid coregistration (warping) of a mean functional image onto the individual anatomies.

Fixed effect individual analyses were performed for each condition in each monkey, with a level of significance set at p<0.05 corrected for multiple comparisons (FWE, t>4.8, unless stated otherwise).

To define the ***preferential near space network***, we contrasted the cortical activations obtained by the stimulation of near space by a small object to those obtained by the stimulation of far space by a large object of the same apparent size as the small object in the near space and was additionally masked by the activations obtained by the stimulation of near space by a small object contrasted by fixation condition (inclusive ‘near space stimulation vs. fixation’ mask, p=0.05 corrected for multiple comparisons, FWE, see Figure 1D, (1)). To define the ***selective near space network***, the above preferential near space network was additionally masked by the activations obtained by far space stimulations (exclusive ‘far space stimulation vs. fixation’ mask, p=0.05 corrected for multiple comparisons, FWE, see Figure 1D, (2)). To define the ***preferential far space network***, we contrasted the cortical activations obtained by the stimulation of far space by a large object of the same apparent size as the small object in the near space to those obtained by the stimulation of near space by a small object and was additionally masked by the activations obtained by the stimulation of far space by a large object contrasted by fixation condition (inclusive ‘far space stimulation vs. fixation’ mask, p=0.05 corrected for multiple comparisons, FWE, see Figure 1D, (3)). To define the ***selective far space network***, the above preferential far space network was additionally masked by the activations obtained by near space stimulations (exclusive ‘near space stimulation vs. fixation’ mask, p=0.05 corrected for multiple comparisons, FWE, see Figure 1D, (4)).

To define the ***preferential encoding of the large cube in the far space network***, we contrasted the cortical activations obtained by the stimulation of far space by a large object to those obtained by the stimulation of far space by a small object and was additionally masked by the activations obtained by the stimulation of far space by a large object contrasted by fixation condition (inclusive ‘Large far space stimulation vs. fixation’ mask, p=0.05 corrected for multiple comparisons, FWE, see Figure 1D, (5)). To define the ***selective encoding of the large cube in the far space network***, the above preferential encoding of the large cube in the far space network was additionally masked by the activations obtained by the stimulation of far space by a small object (exclusive ‘Small far space stimulation vs. fixation’ mask, p=0.05 corrected for multiple comparisons, FWE, see Figure 1D, (6)). To define the ***specific looming toward near space network***, we contrasted the cortical activations obtained by the looming of a small object toward near space to those obtained by the stimulation of near space by a small object and was additionally masked by the activations obtained by the looming of a small object toward near space contrasted by fixation condition (inclusive ‘looming toward near space vs. fixation’ mask, p=0.05 corrected for multiple comparisons, FWE, Figure 1D, (7)) and masked by the activations obtained by near space stimulations (exclusive ‘near space stimulation vs. fixation’ mask, p=0.05 corrected for multiple comparisons, FWE, Figure 1D, (8)).

In all analyses, realignment parameters, as well as eye movement traces, were included as covariates of no interest to remove eye movement and brain motion artifacts. When coordinates are provided, they are expressed with respect to the anterior commissure. Results are displayed on flattened maps obtained with Caret, for each monkey (Van Essen et al. 2001; http://www.nitrc.org/projects/caret/). The results were consistent in the two animals for all the discussed cortical regions (i.e. activations observed in at least 3 out of 4 hemispheres) and do not reveal blink (Guipponi et al., 2015b) or saccade-related activations (see results). Thus, figures 6, 7 and 8 correspond to a group analysis, so as to simplify the presentation of the results. In this case, fixed effect group analyses were performed for each sensory modality and for conjunction analyses with a level of significance set at p<0.05 corrected for multiple comparisons (FWE, t>4.8) and projected onto the anatomy of monkey MT. In order to have an unbiased group analysis, we selected the 15 best runs of MZ (in term of fixation performance), so as to have the same number of runs for each monkey. The results are then displayed on the flattened and fiducial maps of MT.

Assigning the activations to a specific cortical area was performed on each individual monkey brain using the monkey brain atlases made available on http://scalablebrainatlas.incf.org. These atlases allow mapping specific anatomical coronal sections with several cytoarchitetonic parcellation studies. We used the Lewis and Van Essen (2000) and the Paxinos Rhesus Monkey (2000) atlases.

### Regions of interest

We performed regions of interest (ROI) analyses using MarsBar toolbox (Brett et al., 2002), based on the fixed effects individual analyses results. We defined geometric cubic ROIs (2x2x2 mm) centered on the local maximum t-score based on one of the activations obtained for each contrast at FWE-corrected level (*t*-scores>4.8). This ROI analysis only serves to illustrate how PSC changed, within each functional network, as a function of the type of visual stimulation: the specific near space activations, the preference near activations, the unselective near-far space activations, the preference far activations and the specific far space activations. It is thus fully redundant with the flat map analysis. The percent of signal change (PSC) are extracted for each ROI for all the runs using SPM8 and the MarsBar toolbox. The significance of these PSCs across all runs was assessed using a one-tailed paired t-test, in Matlab^TM^ (The MathWorks Inc., Natick, MA, USA).

### Eye movements data analysis

During the MRI scanning sessions, the horizontal (X) and vertical (Y) eye position of the monkey was recorded (in degree) from one eye. An example of the time course of eye position is presented in Figure 2A and 2B, for an exemplar run. For each of these exemplar runs, the block structure is indicated. The refined analysis of eye movements on all the runs per monkey demonstrates that, on average, the monkeys succeeded in maintaining their eyes within the fixation window all throughout the run, irrespective of the type of sensory manipulation (Friedman test >0.05, data not shown). The average and standard deviation of microsaccade and saccade duration with respect to the total blocks duration over all sessions are shown in Figure 2C. These differ as a function of stimulation block (Kruskall-wallis: p<0.001 for MZ and p<0.01 for MT). In supplementary figure S2, a precise description of the expected range of eye movements for each type of stimuli if the monkeys had been fixation and tracking the stimulation Objects rather than maintaining vergence onto the fixation spot is provided. This analysis shows that any tracking of the object would have resulted in a significant increase in out of the fixation window epochs, beyond what is observed experimentally.

**Figure 2:**
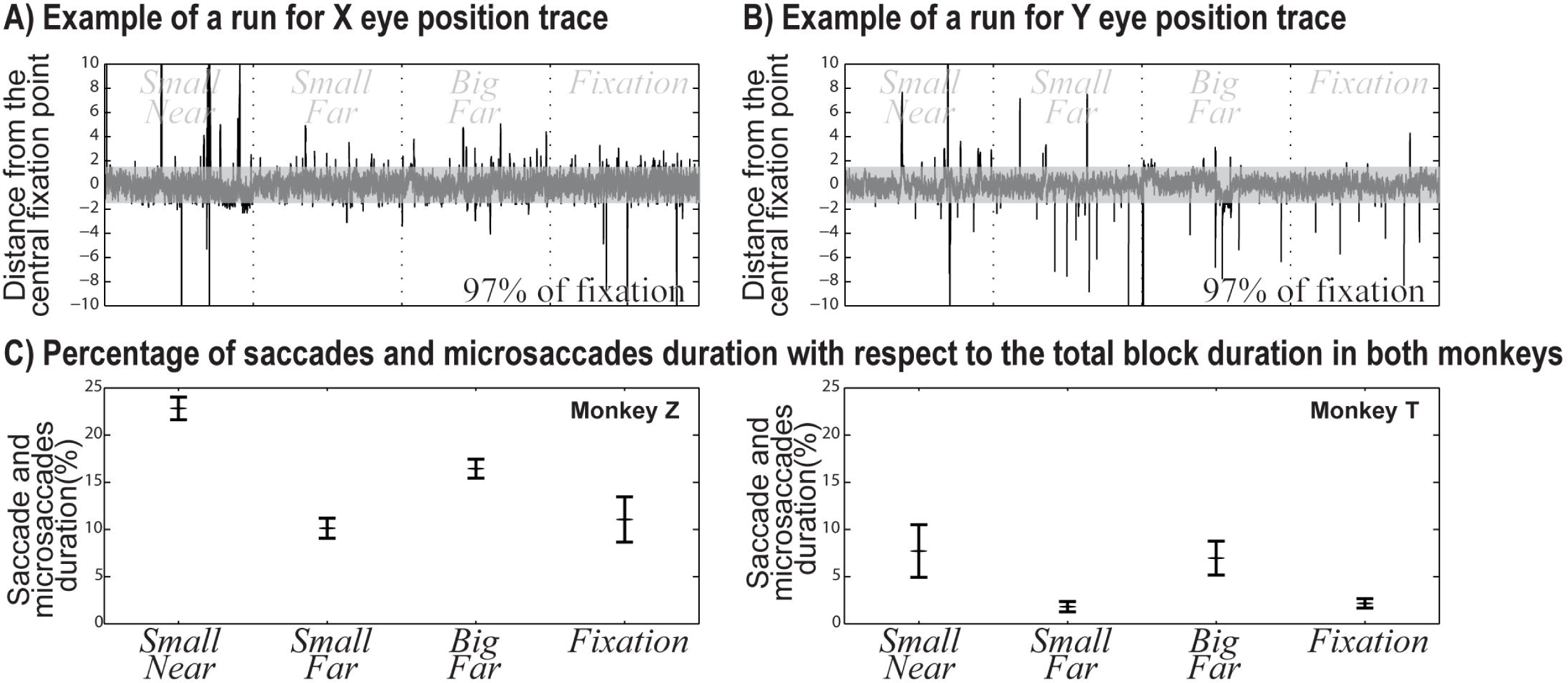
Analysis of eye movements as function of stimulation blocks. 2A) X and 2B) Y eye traces for an exemplar run. 2C) Average and standard deviation of microsaccade and saccade duration with respect to the total blocks duration over all sessions (Kruskall-wallis: p<0.001 for MZ and p<0.01 for MT).

### Signal to noise ratio (SNR)

We calculated the temporal SNR for both monkeys to check for inhomogeneities between the two hemispheres of each animal. The signal is quite homogeneous on the maps (see Supplementary data S3). The apparent asymmetry between the activation strengths on the different contrasts are most probably to the fact that the experimenters were always located on the left side of the monkey, hence the higher right cortical activations.

## RESULTS

Monkeys were exposed, in the same time series, to naturalistic near or far space stimulations (Figure 1), while maintaining their gaze at an intermediate fixation location, so as to keep vergence signals constant all throughout the recording runs. This design allows us to describe the cortical networks involved in near and far space processing in naturalistic conditions. The reported activations in figures 3, 4 and 5 are identified using an individual analysis, with a level of significance set at p<0.05, corrected for multiple comparisons (FWE, t>4.8). The reported activations in figures 6, 7 and 8 are identified using a group analysis, with a level of significance set at p<0.05, corrected for multiple comparisons (FWE, t>4.8). As a result, they reflect the activations that are common to the two monkeys involved in the study.

### Interaction of naturalistic presentations with saccadic behavior

The monkeys are required to fixate a fixation led placed at 90 cm from their face, while being presented with either a small (3cm) cube very close to the face or a large (30cm) cube in the far space. An important question is to characterize the effect of these stimuli onto saccadic behavior and to check whether the monkeys were indeed maintaining vergence onto the fixation led. A detailed analysis of the cumulated microsaccades and saccades duration are shown in Figure 2C. These durations are significantly different following the block stimulation in both monkeys (Kruskall-wallis: p<0.001 for MZ and p<0.01 for MT). Indeed, the percentage of correct fixation changes following the block stimulation: 77.2% (/92.3%) when the small cube stimulated the near space; 89.9% (98.2%) when the small cube stimulated the far space; 83.6% (/93.0%) when the big cub stimulated the far space and 89.0% (97.83%) during fixation only, for monkey Z (/monkey T respectively). In other words, maintaining fixation during the near space stimulation by the small cube was the most challenging. Were the monkeys tracking the near cube? Given the volume of near space that was being explored by the small cube stimulation (Supplementary figure S2, upper row), if the monkey had been tracking the object continuously, the out of the window eye epochs in this condition would have been much larger than in the other conditions. This is an indirect indication that the monkeys did try and suppress eye tracking of the near object during this condition. Nonetheless, could this small systematic behavioral bias affect our functional description of near specific activations? Anticipating on the core of the present work, no specific frontal eye field area or lateral intraparietal area activations are observed during this near space stimulation as compared to the far space stimulation. These two cortical regions are involved in saccadic behavior and their activation is observed during saccadic behavior (Everling and Munoz, 2000; Hutchison et al., 2012; Koyama et al., 2004; see for review Womelsdorf and Everling, 2015).

### Naturalistic near and far space stimulations

Naturalistic near space stimulations with a small cube (Figure 3, yellow upper maps, small near vs. fixation contrast) activated a small extent of the occipital striate and extrastriate areas, the temporal cortex (superior temporal sulcus), the parietal cortex, the prefrontal cortex (arcuate sulcus and posterior and anterior parts of principal sulcus) as well as the orbitofrontal cortex. Far space stimulations with a far cube with the same apparent size as the near small cube (Figure 3, blue middle maps, big far vs. fixation contrast) also activated a widespread cortical network including the entire striate and extrastriate cortex, the temporal cortex, the parietal cortex, small portions of the prefrontal cortex along the arcuate sulcus as well as the orbitofrontal cortex. When far space was stimulated using a cube of the same real size as the small cube used for the near space stimulation (Figure 3, green lower maps, small far vs. fixation contrast), a similar though smaller cortical network was activated. In the following, we identify those cortical regions that are either preferentially or specifically involved in near space and far space processing respectively.

**Figure 3:**
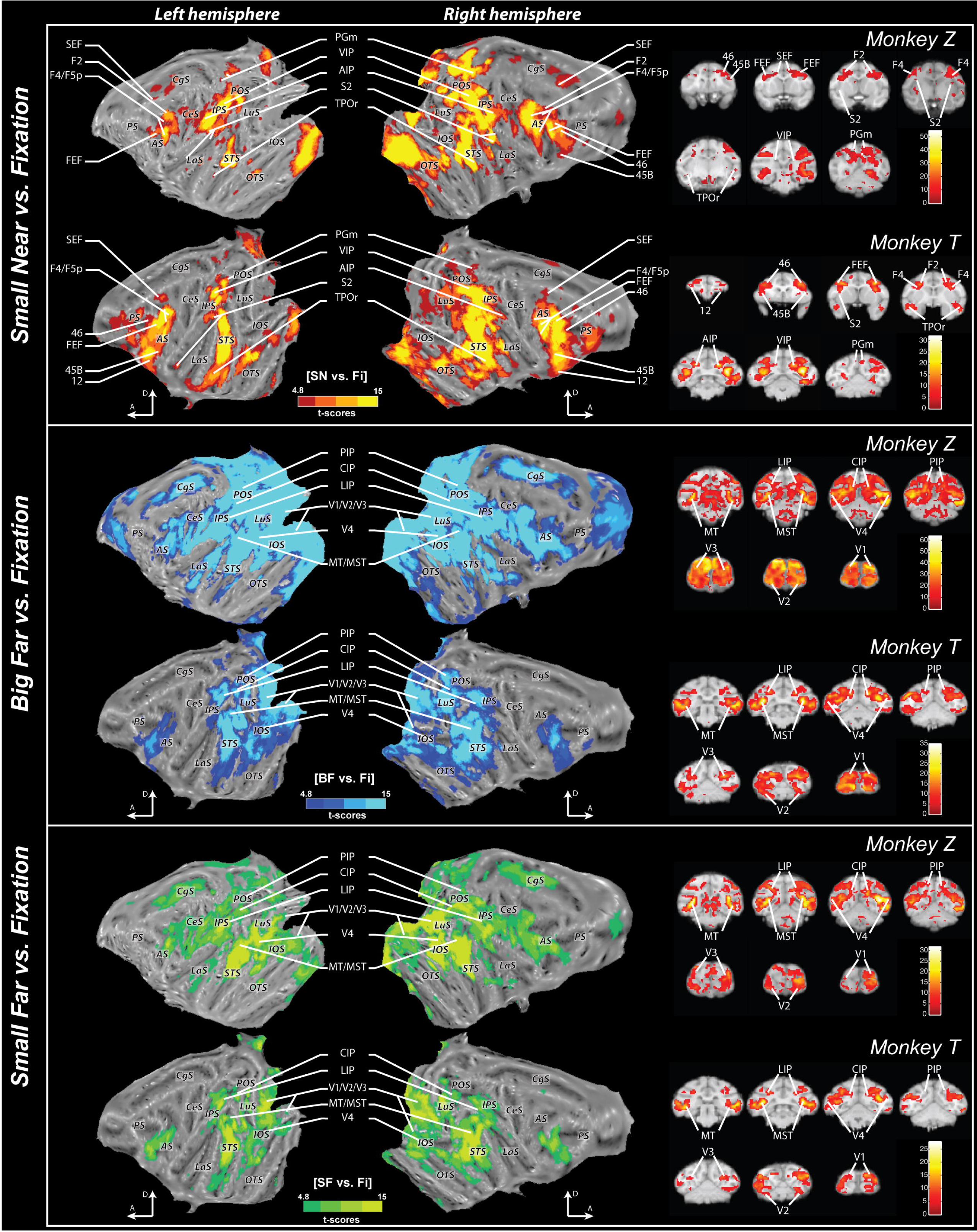
Near space and far space individual analyses. Activations are presented on the flattened representation of the two monkey right and left hemispheres obtained with Caret. The upper part of the figure shows the near space stimulated with the small cube (SN) versus fixation contrast (t scores = 4.8 at p<0.05, FWE-corrected level in the red scale). Middle and lower panels present the far space respectively stimulated with the big (BF) and the small cubes (SF; t scores = 4.8 at p<0.05, FWE-corrected level respectively in the blue and green scales). The right panel represents selected coronal slices showing the activated areas for each contrast in both monkeys. A, Anterior; D, Dorsal; SN: small near; BF: big far; SF: small far. 12, area 12; 45B, area 45B; 46, area 46; AIP, anterior intraparietal area AIP; CIP caudial intraparietal area; F2, premotor area F2; FEF, frontal eye field; LIP, lateral intraparietal area LIP; MT/MST, medial/superior temporal areas MT/MST; F4/F5, premotor areas F4/F5; PGm, medial parietal area PGm; PIP, posterior intraparietal area PIP, S2, somatosensory area 2; SEF: supplementary eye field; TPOr, rostral temporoparietal occipital area TPOr; VIP, ventral intraparietal area VIP; V1/V2/V3/V4, visual areas V1,V2,V3 or V4. Cortical sulci: AS, arcuate sulcus; CgS, cingulate sulcus; CeS, central sulcus; IOS, inferior occipital sulcus; IPS, intraparietal sulcus; LaS, lateral (Sylvian) sulcus; LuS, lunate sulcus; OTS, occipital temporal sulcus; POS, parieto-occipital sulcus; PS, principal sulcus; STS, superior temporal sulcus.

In this Figure 3, the near space stimulation vs. fixation activation maps contrasts with the far space stimulation vs. fixation activations maps in that central striate and extra-striate cortex is spared in the former condition. While this can seem extremely surprising (as strong central visual stimulation is expected to reliably activate these regions), this observation that is reproduced in both monkeys corresponds to what one would expect if the monkeys were fixating the fixation led and suppressing visual information away from the fixation plane. Indeed, if one assumes an eye-to-eye distance of 2.6 cm, fixation vergence angle would be around 1.65° and near cube stimulation would correspond to crossed visual binocular disparities of 8°. V1 neurons are reported to be selective for absolute binocular disparities below 1°, thus possibly accounting for our observations in this respect.

### Distinctions within the near space cortical network

In the following, we define three different functional near space networks: a restricted network selectively encoding near space; a larger network preferentially encoding near space with respect to far space and an even larger network encoding near space irrespectively of any coding for far space. The larger ***non-selective near space network*** corresponds to the one identified in Figure 3 (yellow upper maps, represented in the Figure 4 as a red contour) and discussed in the previous section. The ***preferential near space network*** (Figure 1D (1), Figure 4 A and C, dark red, Figure 8, dark red), activates bilateral cortical regions the contribution of which is statistically higher for near space than for far space. These include parietal areas: the posterior intraparietal sulcus (IPS: ventral intraparietal area VIP, the posterior medial intraparietal area MIP) as well as its anterior most tip (possibly anterior intraparietal area AIP), the medial parietal cortex (area PGm) and the parietal opercular region area 7op; temporal areas: the rostral temporoparietal occipital area TPOr in the medial mid-to-anterior bank of the superior temporal sulcus, the intraparietal sulcus associated area IPa, the inferior temporal area TEAa-m, the dorsal portion of the subdivision TE1-3; insular regions: the parainsular cortex PI; somatosensory area SII within the medial bank of the lateral sulcus; prefrontal and premotor regions: dorsal premotor cortex F2, premotor area 4C or F4/F5 including premotor zone PMZ, the supplementary eye field, the frontal eye fields (area 8a as well as 8ac), prefrontal area 46p, prefrontal area 45B; frontal regions: the posterior orbitofrontal area 12.

**Figure 4:**
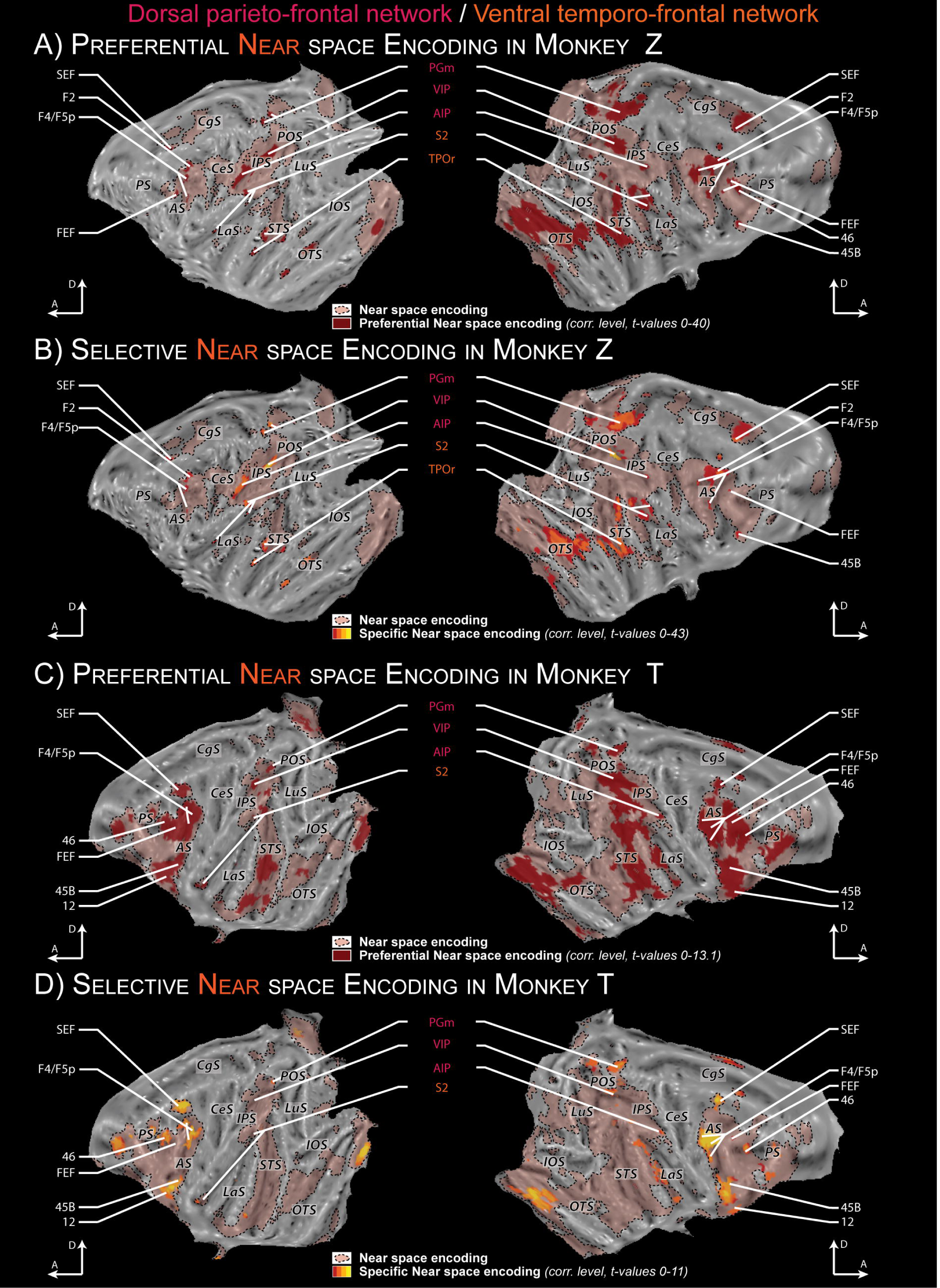
Non-selective, preferential and specific near space networks. Activations are presented on the flattened maps of individual monkeys. Only the key activations identified in three hemispheres out of four are labelled. A and C) Non-selective and preferential near space encoding for monkey Z (A) and T (C). Preferential near space coding corresponds to the cortical regions whose activations are higher for the small cube in near space than for the large cube in far space (t scores = 4.8 at p<0.05, FWE-corrected level in the dark red). B and D) Non-selective and specific near space encoding for monkey Z (B) and T (D). Specific near space encoding corresponds to the cortical regions which are activated by the small near cube but not for the large far cube (red to yellow scale, exclusive mask for far space versus fixation baseline applied at FWE-corrected level p<0.05). The outer red contours correspond to non-selective near space encoding (near space versus fixation, as in figure 3). For other conventions, see Figure 3.

The ***selective near space network*** (Figure 1D (2), Figure 4 B and D, red to yellow color scale,Figure 8, colored to yellow color scale, *t* scores = 4.8 and above, FWE-corrected level), exclusively involved in near space processing activates discrete bilateral regions within the majority of the cortical areas highlighted by the previous contrast.

### Distinctions within the far space cortical network

Likewise, we define three different functional far space networks: a restricted network selectively encoding far space; a larger network preferentially encoding far space with respect to near space and an even larger network encoding far space irrespectively of any coding for near space. The larger ***non-selective far space network*** corresponds to the one identified in Figure 3 (blue middle maps, represented in the Figure 5 as a blue contour) and discussed above. The ***preferential far space network*** (Figure 1D (3), Figure 5 A and B: dark blue, Figure 8, dark blue), activates bilateral cortical regions the contribution of which is statistically higher for far space than for near space. A strong inter-individual variability can be observed, these activations being larger in Monkey Z than in Monkey T. Differences in global activation strength are often observed when performing single subject fMRI analyses. In the present context, an alternative hypothesis can be put forward. Monkey Z is a small female while monkey T is a large dominant male. The larger activations observed in monkey Z could indicate higher overall activations to the presented naturalistic stimuli because of their higher relative size with respect to body size, as compared to monkey T. This would need to be confirmed experimentally. In the following, only the common activations are described. These encompass the entire visual striate and extrastriate cortex: areas V1, V2, V3, V3A and V4; they also include parietal cortical regions: the posterior intraparietal area PIP, the lateral intraparietal area LIP, the medial parietal convexity (area 5v), the lateral parietal convexity (area 7a, 7ab and 7b), as well as the posterior most part of the intraparietal sulcus (caudial intraparietal area CIP, parietal reach region, PRR), these activities extending towards the parieto-occipital cortex (including areas V6A and V6); temporal cortex: medial and superior temporal area MT and MST.

**Figure 5:**
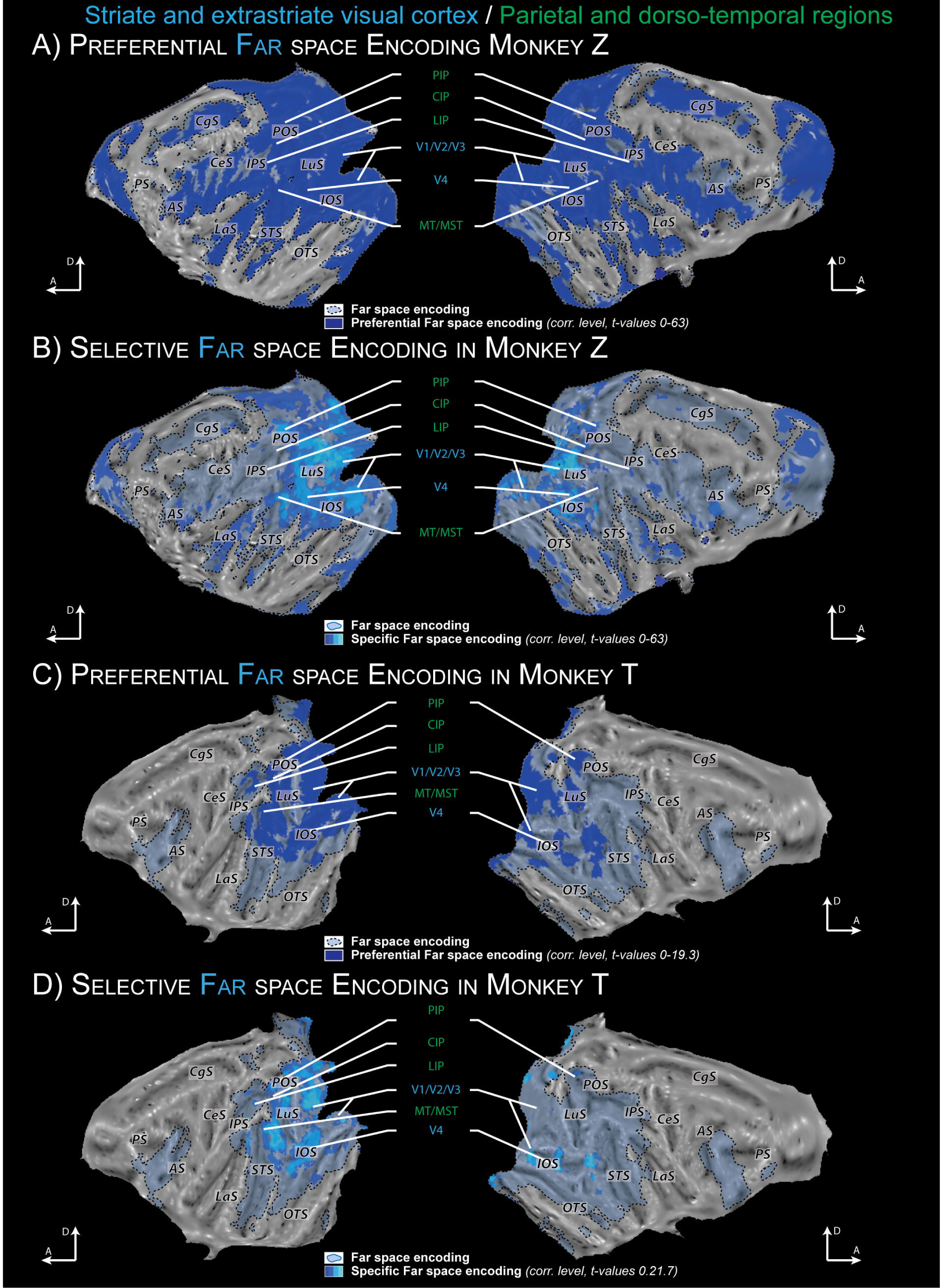
Non-selective, preferential and specific far space networks. Activations are presented on the flattened maps of individual monkeys. Only the key activations identified in three hemispheres out of four are labelled. A and C) Non-selective and preferential far space encoding for monkey Z (A) and T (C). Preferential far space coding corresponds to the cortical regions whose activations are higher for the large cube in far space than for the small cube in near space (t scores = 4.8 at p<0.05, FWE-corrected level in the dark blue). B and D) Non-selective and specific far space encoding for monkey Z (B) and T (D). Specific far space encoding corresponds to the cortical regions which are activated by the large far cube but not for the small near cube (dark blue to light blue color scale, exclusive mask for near space versus fixation baseline applied at FWE-corrected level p<0.05). The outer blue contours correspond to non-selective far space encoding (far space versus fixation, as in figure 2). For other conventions, see Figure 3.

When this contrast is additionally masked by the activations obtained by near space stimulations, thus defining the ***selective far space network***, a network that is selectively involved in far space processing can be identified (Figure 1D (4), Figure 5 C and D, dark blue to light blue color scale, Figure 8: dark blue to light blue color scale, *t* scores = 4.8 and above, FWE-corrected level). This analysis describes discrete bilateral regions essentially in occipital and temporal areas highlighted by the previous contrast and few parietal areas.

### Far space cortical network modulation by object size

While the small cube presented in near space had the same apparent size as the far cube presented in far space, these two objects had very different physical sizes (3x3x3cm^3^versus 30x30x30cm^3^). As a result, part of the far or near space network specificities described above could have been due to this objective size difference (as opposed to an apparent size difference). In order to address this issue, we now compare the cortical activations obtained when stimulating far space with either a small cube or a large cube. For the sake of space, a group analysis is performed rather than a single subject analysis. This group analysis captures the common activations already described at the single individual level in figures 3, 4 and 5. No activations are observed with the *small object in far space* versus *large object in far space* contrast, indicating that all the cortical regions that are involved in processing the small object in far space also contribute to the processing of the large object in far space. The inverse contrast reveals a large cortical network (Figure 1D (5), Figure 6, dark blue) mostly identical to that revealed by the *large cube in far space* versus *fixation* contrast (Figure 3, middle blue panels). The network that is selective to the large cube in far space processing as compared to the processing of smaller objects in far space (Figure 1D (6), Figure 6, dark blue to light blue color scale, *t* scores = 4.8 and above, FWE-corrected level) includes large sectors of the visual striate and extrastriate cortex, mostly coinciding with the peripheral visual field representation (see Figure 4 in Guipponi et al., 2015 for an identification of these representations on this same group of subjects), the parieto-occipital cortex, the posterior parietal cortex, the right medial parietal cortex the anterior part of the superior temporal sulcus as well as a large extent of the prefrontal cortex, the cingulate cortex, the orbitofrontal cortex and the frontal pole.

**Figure 6:**
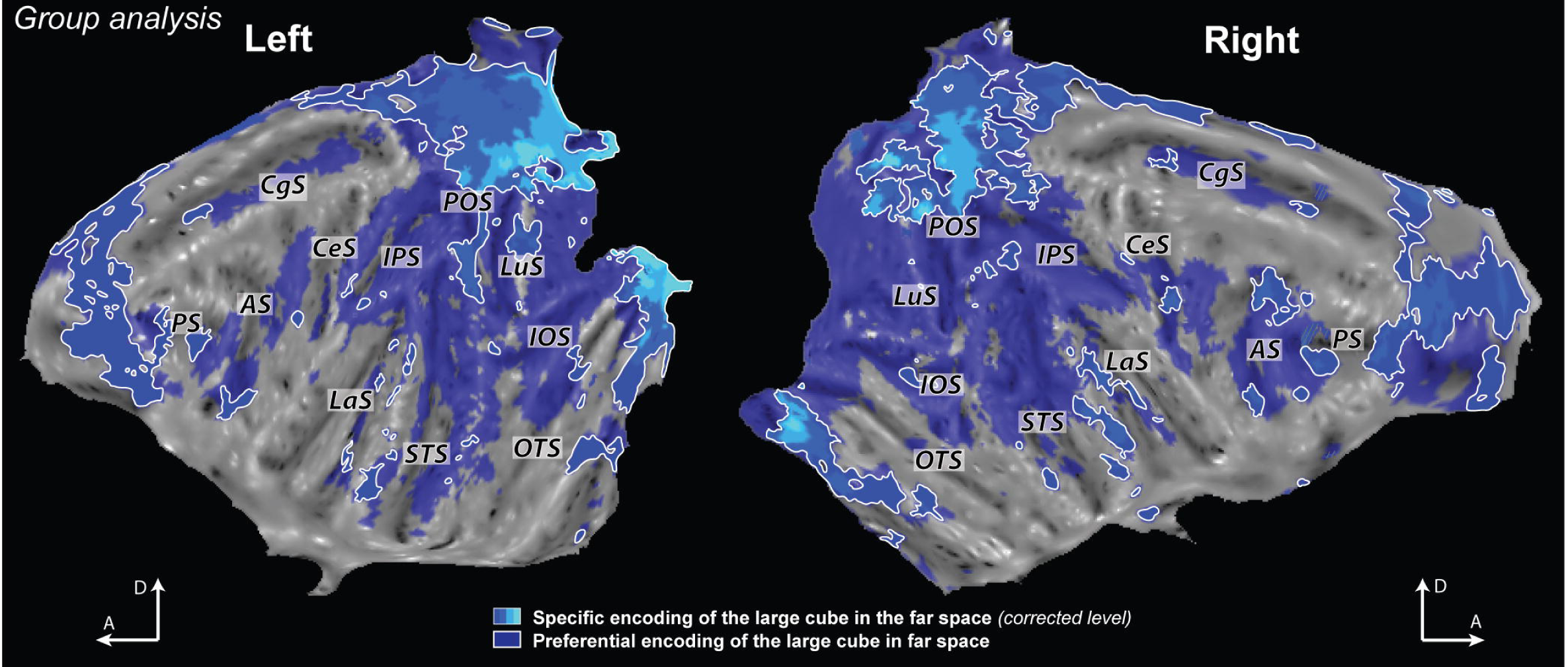
Preferential and specific encoding of the large cube in the far space. Activations are presented on the flattened maps of the reference monkey cortex (group analysis). Preferential encoding of the large cube in the far space corresponds to the cortical regions whose activations are higher for the big cube in far space than for the small cube in far space (t scores = 4.8 at p<0.05, FWE-corrected level in the dark blue). Specific encoding of the large cube in the far space corresponds to the cortical regions which are activated by the big cube but not for the small far cube in the far space (dark blue to light blue color scale, exclusive mask for the big cube in far space versus fixation baseline applied at FWE-corrected level p<0.05). For other conventions, see Figure 3.

### Functional activations in response to looming and recession in near and far space

The physical approach of the cubes into far or near space and their recession back behind the curtains was controlled by the experimenters following a precise schedule indicated to them on a computer screen. The onsets of these displacements were logged together with all other task events. Here, we focus on the activations observed during these dynamic phases of the task (Figure 7). For the sake of space, a group analysis is performed rather than a single subject analysis. *The approach of the small cube into near space* produces widespread activations (Figure 7A). This network includes the orbitofrontal cortex (12, 46p), prefrontal and premotor cortex (FEF, F4, F5), parietal cortex (VIP, PIP, 7a, 7b, 7ab), temporal cortex (MT, MST, TPOr, IPa) and visual areas (V1, V2, V3). This network is mostly included in the near space network (Figure 7A, black contours, Figure 3, top panel), though a heterogeneity can be noted. Some cortical regions modulated by near space stimulations are not activated by intrusion into near space, mostly along the ventral visual stream (Figure 7A, uncolored cortex within the black contours). Other cortical regions modulated by near space stimulations are equally activated by intrusion into near space and near space stimulations (Figure 7A, colored activations within the black contours). Importantly, not all the regions that are activated by intrusion into peripersonal space are also activated by the sustained presence of a stimulus in near space (Figure 1D (7), Figure 7A, white contours outside the black contours). *The approach of the big cube into far space* produced very similar activation patterns to those observed during the approach of the small cube into near space, to the notable exception of the orbitofrontal cortex, possibly indicating an emotional component to intrusion into near space (Figure 7E). In contrast, *the approach of the small cube within far space* produced more restricted activations, mostly in the intraparietal sulcus (IPS), in the superior temporal sulcus (STS) as well as in the striate and extrastriate cortex (Figure 7C).

**Figure 7:**
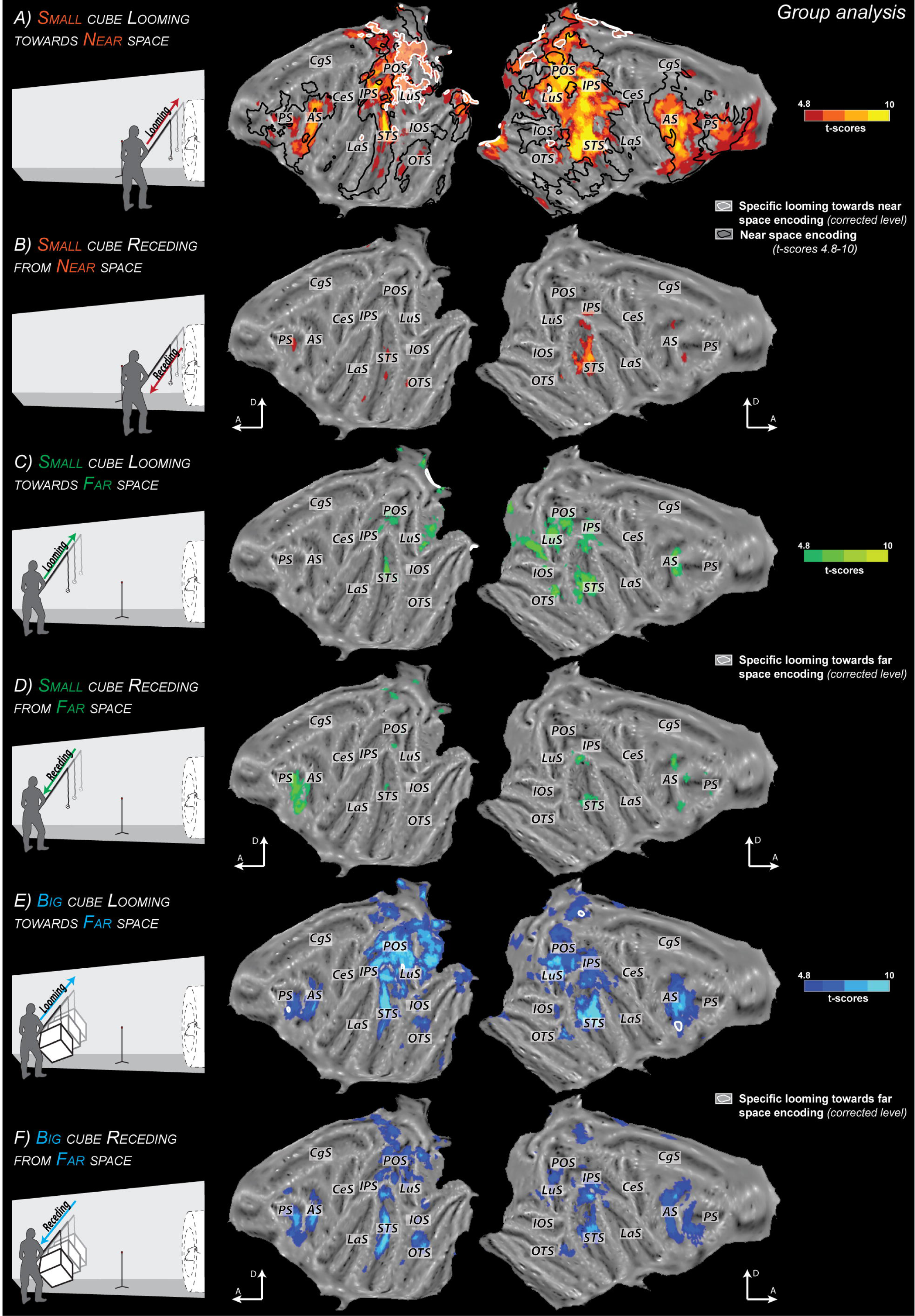
Looming stimuli activate both Near and Far space network. Activations are presented on the flattened maps of the reference monkey cortex (group analysis). A, C and E, this presents the approach of the stimulus respectively towards near space with the small cube, towards far space with the small cube or towards far space with the big cube versus fixation contrast (t scores = 4.8 at p<0.05, FWE-corrected level respectively in the red, green and blue scale). B, D and F, this presents the receding of the stimulus respectively from near space with the small cube, from far space with the small cube or from far space with the big cube versus fixation contrast (t scores = 4.8 at p<0.05, FWE-corrected level respectively in the red, green and blue scale). On the first panel (the approach of the stimulus towards near space with the small cube), the black contours represent the near space encoding (one identified in Figure 3) and the white contours represent the specific encoding of the approach of the small cube in the near space which corresponds to the cortical regions which are activated by the approach of the small cube in the near space but not by the near space encoding (exclusive mask for the near space versus fixation baseline applied at FWE-corrected level p<0.05). For other conventions, see Figure 3.

*When the small cube recedes from near space*, very few activations are observed, circumscribed to the prefrontal area 46 and the superior temporal sulcus STS (Figure 7B). A very similar activation pattern is described for *the recession of the small cube from far space* (Figure 7D). Last, activations are more widespread for *the large cube receding from far space*, quite close to those observed for the approach of this stimulus into far space (Figure 5F). This contrasts with the difference observed between the looming and recession of the small cube in near space, and support the idea that the representation of object movement vectors in the cortex differ depending on whether movement takes place in near or in far space.

### Near to far coding gradient in the arcuate (AS) and intraparietal sulcus (IPS)

Intraparietal and periarcuate regions have been shown to play a key role in space representation and space representation for action. In this section, we focus on these two regions (Figure 8). Again, for the sake of space, a group analysis is performed rather than a single subject analysis. This group analysis captures the common activations already described at the single individual level in figures 3, 4 and 5. In the *Arcuate sulcus*, near space is specifically encoded by areas 46p, 12, F4, F5 and SEF whereas specific far space is overall represented at the inferior tip of the AS and along the gyrus posterior to the AS. In the *Intraparietal sulcus*, near space is specifically encoded by the VIP area and PGm area whereas far space is specifically encoded by the areas 5v and PIP.

**Figure 8:**
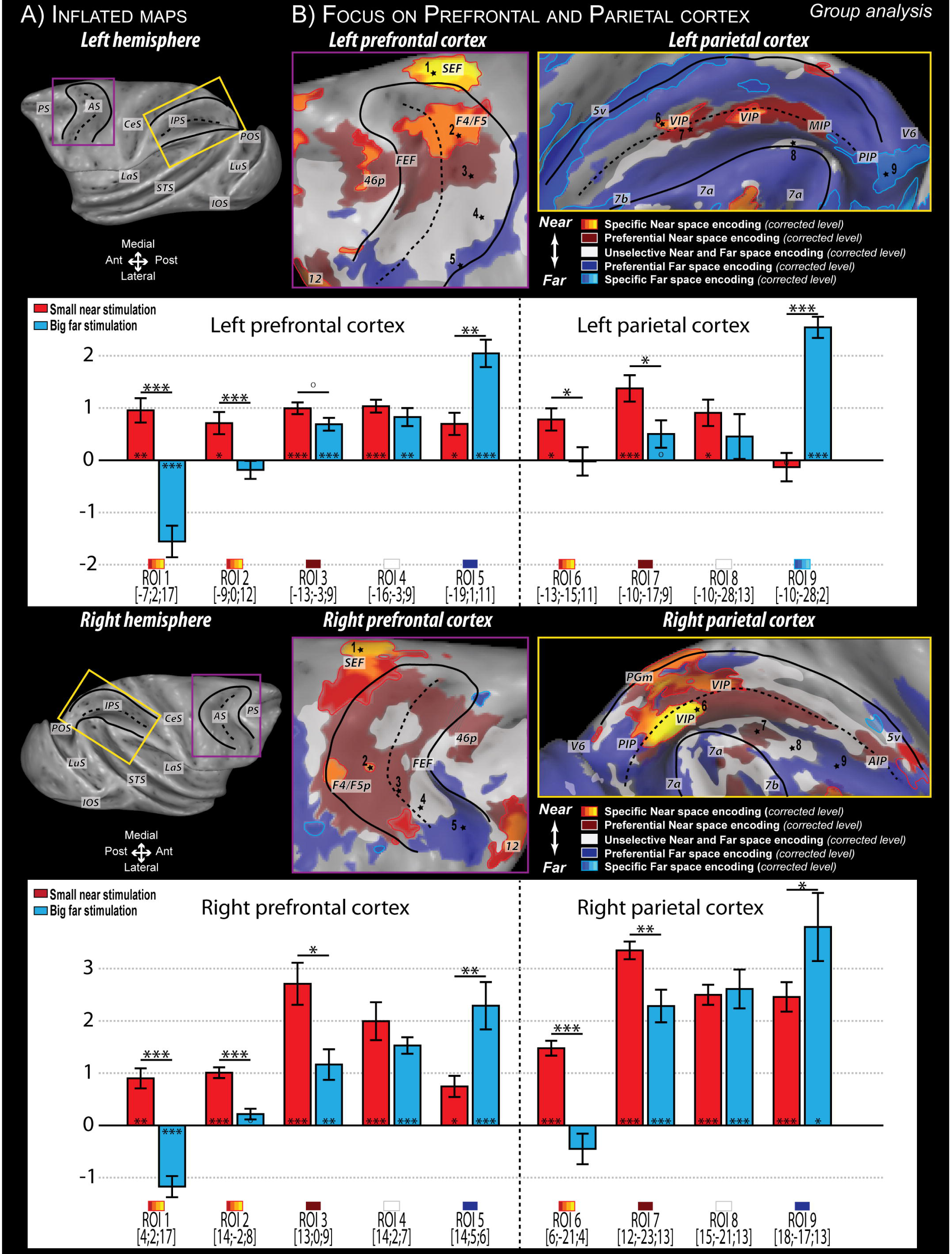
Near to Far coding gradient in the arcuate sulcus (AS) and the intraparietal sulcus (IPS). A) Inflated representation of the cortex, (left and right hemisphere of the reference monkey cortex). The purple inset corresponds to the AS and the yellow inset corresponds to the IPS as represented in B). Black solid lines indicate the limit between the convexity and the banks of the IPS and AS; black dashed lines indicated the bottom of the sulcus. B) Prefrontal (purple inset) and parietal (yellow inset) activations of near and far space networks (group analysis superimposing the activations described in figure 4 and 5) on the same inflated maps, for the left and right hemispheres. Red to yellow color scale: selective near space coding; Red: preferential coding for near space; White: unselective coding for near and far space; Blue: preferential coding for far space; Dark blue to light blue color scale: selective near space coding. Gray regions correspond to regions activated neither by the large stimulus in far space nor by the small stimulus in near space. 5v, ventral area 5v; 7a,7b/7ab, areas 7a, 7b or 7ab: MIP, medial intraparietal area MIP, V6: cortical visual area V6. For other conventions, see Figure 3, 4 and 5. Five regions of interest were defined along the AS and four along the IPS, and percentage of signal change (PSC) within each of these ROIs was extracted for each contrast of specific near space, preferential near space, unselective near and far space, preferential far space and specific far space. Below each inflated map, the PSCs are plotted for the stimulation of the small cube in the near space (orange bars) and for the stimulation of the big cube on the far space (blue bars). The specific location of each ROI as well as the map they belong to is indicated at the bottom of the figure. Statistical significance is represented as follow: *p<0.05; **p<0.01; ***<0.001 (paired t-tests with Bonferroni correction); **O** p<0.05 (paired t-tests, uncorrected).

Different regions of interest were defined along the AS and the IPS, and the percentage of signal change (PSC) within each of these ROIs was extracted for each contrast of specific near space, preferential near space, unselective near and far space, preferential far space or specific far space, Figure 8, histograms). Overall, this analysis confirms the existence, along the AS, bilaterally, of a progressive increase in the PSC to far stimuli (central blue bars) from ROIs 1 to 5 (176% of increase for the left and 151% for the right), together with a quite stable PSC for near stimuli (central red bars). A similar progressive increase in the PSC to far stimuli can also be observed in the IPS, bilaterally (ROIs 6 to 9, central blue bars, 101% for the left and 112% for the right), together with a stable PSC for near stimuli (central red bars). Confirming the whole brain contrast analysis, we also note deactivations or non-significant activations during the stimulation of far space by a larger object in the specific near space contrast (ROIs 1, 2 and 6, bilaterally) and the preferential near space contrast (ROIs 3 and 7, bilaterally). Likewise, the PSC during the stimulation of far space by a larger object are significantly higher than the PSC during the stimulation of near space in the specific and preference contrasts (ROIs 5 and 9, bilaterally). In the unselective near and far space contrast, the PSCs during the stimulation of near space and far space are not significantly different (ROIs 4 and 8, bilaterally).

## DISCUSSION

In the following, we discuss in the identified primate networks associated with either near or far space stimulation with naturalistic dynamic objects in the light of the related literature.

### Near space specific cortical network

The near space specific cortical network we describe here is surprisingly large. It involves multisensory visuo-tactile cortical regions whose neurons have already been described to encode nearby objects relative to the body and subserve peripersonal space representation, namely parietal area VIP (Duhamel et al., 1997, 1998; Avillac et al., 2005; Schlack et al., 2005; Bremmer et al., 2013; Guipponi et al., 2013) and premotor area F4 (Cléry et al., 2015b; Fogassi et al., 1996; Graziano et al., 1994, 1997, 1999; Matelli and Luppino, 2001; Rizzolatti et al., 1981). These two regions have anatomical connections and functional homologies. This VIP-F4 network processes all the necessary information to bind together the localization of objects around our body with actions towards these objects and to define a safety body margin contributing to the definition of self with respect to the external world (Graziano and Cooke, 2006; Brozzoli et al., 2013, 2014; Chen et al., 2014; Cléry et al., 2015b). In our study, area VIP is sensitive to dynamic stimuli in near space whereas a previous study did not identify a sensitivity to the 3-dimensional structure of static stimuli in VIP (Durand et al., 2007). In a more recent study and in agreement with our results, Van Dromme et al., (2016) show that VIP is involved in depth-structure processing. On the other hand, the parietal area AIP and premotor area F5, activated here by near space stimulations, have been shown to be essential for grasping and reaching processing (Fogassi et al., 2001; Gallese et al., 1994; Murata et al., 2000, 1997; Sakata and Taira, 1994). These two areas share also strong anatomical connections and functional homologies and have consequently been proposed to form a second parieto-premotor network dedicated for action (Gallese et al., 1996; Iriki et al., 1996; Matelli and Luppino, 2001; Rizzolatti and Luppino, 2001; Rizzolatti and Matelli, 2003; see for discussion Cléry et al., 2015b)(see for discussion Cléry et al., 2015b; Gallese et al., 1996; Iriki et al., 1996; Matelli and Luppino, 2001; Rizzolatti and Luppino, 2001; Rizzolatti and Matelli, 2003).

We also identify cortical areas whose contribution to near space processing has been partly overlooked. We describe the involvement of dorsal premotor regions, in agreement with the description, in patients with dorsolateral prefrontal cortex damage, of a near space specific neglect (Aimola et al., 2012). We also describe the involvement of posterior, fundal and medial parietal areas, in agreement with the description of a near space neglect in patients with posterior parietal lesions (Halligan and Marshall, 1991). Near space activations are also observed within the fundus of the STS and on the inferior temporal convexity, suggesting a specific processing of near objects feature and identity within the ventral visual stream.

Near space specific activations are also observed in area SII, possibly revealing a general “attention-to-touch” process due to the anticipation of tactile stimulation to the body. This could also actuate the strong functional link between near space processing and the somatosensory representation of self. While previous studies have mostly assumed that this link is subserved by associative multisensory visuo-tactile areas (Blanke, 2012; Makin et al., 2008), the present observations suggest that low level sensory areas might also be involved in the representation of space at the frontier of self. Close to the face, the moving cube can be viewed as a potentially dangerous object. This could possibly also account for the observed orbitofrontal activations (area 12, Murray and Izquierdo 2007).

Overall, the non-human near space specific cortical network we describe here has major specificities as compared to the analog human network. We essentially describe bilateral cortical regions, while in humans, only the left dorsal occipital cortex, the left intraparietal and the left ventral premotor appear to be involved in near space processing (Aimola et al., 2012; Weiss et al., 2000). This near space network areas also appears larger in the monkeys than in humans. This could be due to the fact that we stimulated near space at 15cm away from the subject while Weiss et al. (2000)used stimulations at 70cm. Alternatively, this could be a genuine interspecies difference.

### Far space specific cortical network

The observed monkey far space specific cortical network is also very extended, involving large portions of the occipital cortex, posterior temporal and superior temporal regions, similar to what is seen in humans (Weiss et al. (2000). This is in agreement with the far space neglect following a temporal hematoma (Vuilleumier et al., 1998).

In our study, we show a clear far space preference coding in V6A. A human study (Quinlan and Culham, 2007) shows a near preference coding in dPOS. This is suprising given the high degree of similarity between the human superior parieto-occipital cortex (sPOC) and the macaque parieto-occipital area including areas V6 and V6A (Galletti et al., 1999, 2005). This discrepancy can be explained by important experimental differences. Whereas we use naturalistic and dynamic stimuli in near or far space while monkeys fixated an intermediate position, in Quinlan and Culham (2007), participants fixated in near vs. far space with no other visual stimulation. Therefore, only oculomotor signals of vergence eye position and ocular accommodation are manipulated by the task design.

Last, Rizzolatti et al. (1983) describe a more pronounced hemineglect in far than in near space following prearcuate area 8 ablations whereas we describe a preferential though not specific coding for near space in this region. This discrepancy could reflect a task dependence of far and near space processing as described in humans, in active oculomotor or reaching contexts (Aimola et al., 2012; Keller et al., 2005).

### Object size effects

All the cortical regions involved in processing the small object in far space also contribute to the processing of the large object in far space. The reverse is not true. At the neuronal level, this possibly suggests a multiplexing of object real-size and location in far space relative to the subject, as a big object is not expected to have the same valence as a small object in far space. Size coding might further be normalized with respected to the actual body size of the subject, big objects being possibly more dangerous for small individuals than for larger individuals.

Human studies show that visual objects may be mapped along the ventral occipito-temporal cortex according to their real-world size, reflecting the visual or functional properties associated with small versus big real-world objects (Konkle and Oliva, 2012) but also abstract and conceptual size representation (Gabay et al., 2016). It would be highly interesting to test different objects with real-world size well known for monkey to further characterize the interaction between real object-size and object distance from the subject.

### Looming stimuli and peripersonal space

The looming of the small cube in near space results in large activations involving both the near and far space networks. Such looming stimuli have been described to trigger stereotyped defense responses and enhance reaction times or sensitivity to a second stimulus (Schiff et al., 1962; Ball and Tronick, 1971; Vagnoni et al., 2012; Canzoneri et al., 2012; Kandula et al., 2015; Cléry et al., 2015a) including nociceptive stimuli (De Paepe et al., 2016). Recently, we have described that this type of dynamic visual stimuli activate a parieto-frontal network highly overlapping with the one described here (Cléry et al., 2015b, 2017)(Cléry et al., 2017, 2015b). This functional overlap between a network encoding the presence of a stimulus within peripersonal space at the same time as intrusion into peripersonal space reinforces the view that this network encodes visual stimuli in relation with the margin of self and their possible tactile consequences on the body.

### Substrates for dynamic space representation

The description of specific networks for near and far space processing should not have us overlook the fact that large cortical regions contribute to the processing of both far and near objects, though often favoring one over the other as in LIP, 7a or 7b. Bimodal visuo-tactile neurons have been described in area 7b with large receptive fields over the arm, leg, chest or whole body (Leinonen et al., 1979). Lesions of this region induce a neglect in peripersonal space (Matelli et al., 1984), suggesting that area 7b is involved in the perception of near space and in the organization of movements towards stimuli presented in peripersonal space. However, we found a privileged coding of far space for this area, calling for reassessment of its functional role in relation with space processing.

The description of large cortical regions having either a preference for near or far space processing call for reappraising far space or near space specificity. Indeed, the alternative view we would like to suggest is that of a continuous encoding of relative distance to the body, in the form of a far-to-near gradient. In this context, far or near space specific regions represent the extreme points of this continuum. The idea of such a continuum is supported by the fact that no abrupt change in visuo-spatial neglect can be seen between near and far space (Cowey et al., 1999). Another indirect evidence can be found in a recent non-human fMRI study (Joly et al., 2009), which describes disparity-related signals in far space in area F5a, close to periarcuate far space activation in our Figure 8. It is important to note that the existence of such a cortical far-to-near gradient in space representation does not preclude the existence of a physically delineable peripersonal space (Berti and Frassinetti 2000; Macaluso and Maravita 2010; Farnè et al. 2005; Ladavas and Serino 2008).

Our results open the way to study how these two networks dynamically interact during action planning, tool use or as a function of the emotional or social contexts as some studies show that these processes are dynamics (Markman and Brendl, 2005; Bassolino et al., 2010; Brozzoli et al., 2010; Lourenco and Longo, 2009; Lourenco et al., 2011; Valdés-Conroy et al., 2012; Teneggi et al., 2013).

## Acknowledgments

J.C. was funded by the Fondation pour la Recherche Médicale and by the Fondation Berthe-Fouassier. O.G. was funded by the French education ministry. S.BH was funded by the French Agence nationale de la recherche (Grant #ANR-05-JCJC-0230-01).

---

**Figure S1:**
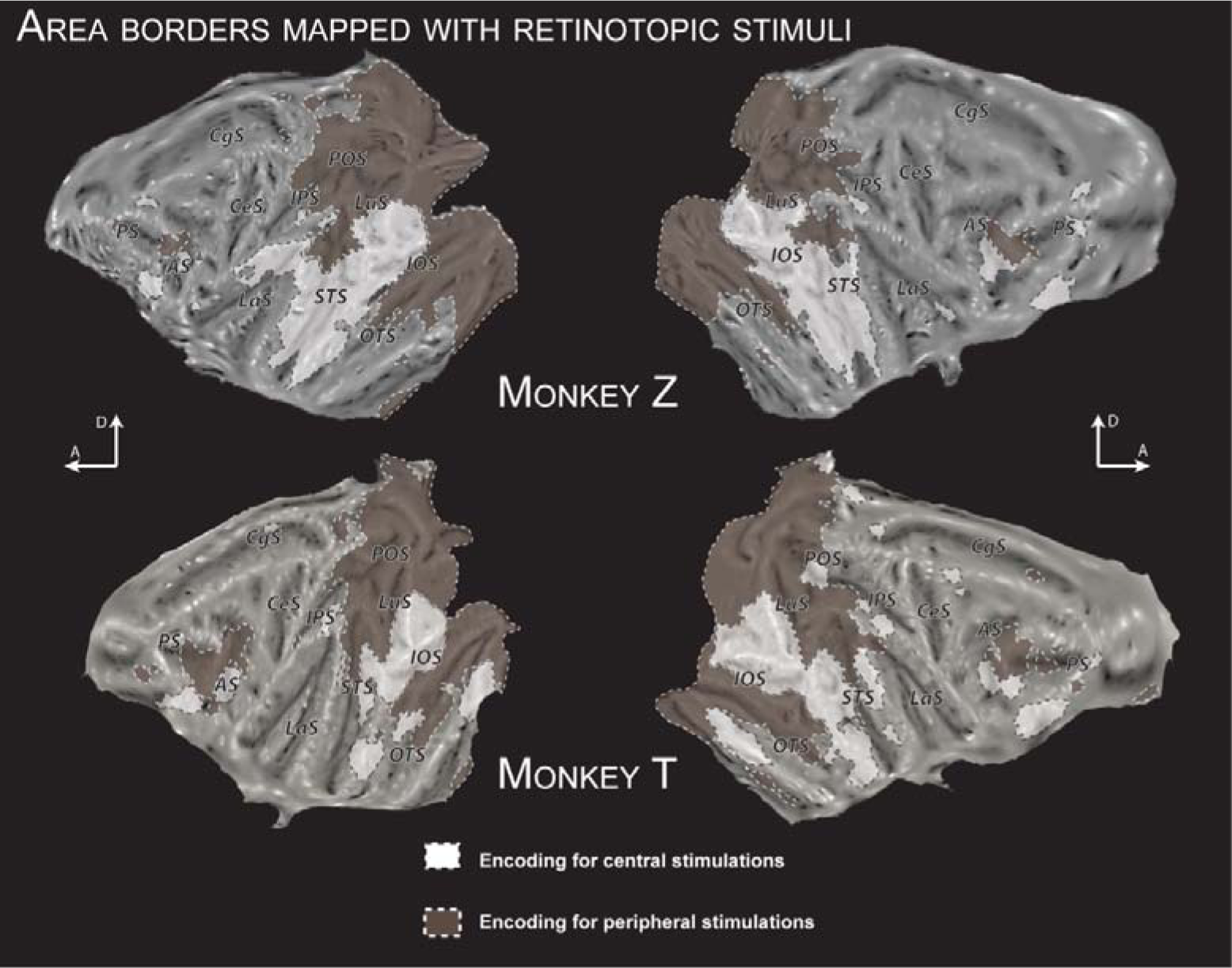
Individual whole brain maps of center vs. periphery visual coding from (Guipponi et al., 2015a). Cortical sulci: AS, arcuate sulcus; CgS, cingulate sulcus; CeS, central sulcus; IOS, inferior occipital sulcus; IPS, intraparietal sulcus; LaS, lateral (Sylvian) sulcus; LuS, lunate sulcus; OTS, occipital temporal sulcus; POS, parieto-occipital sulcus; PS, principal sulcus; STS, superior temporal sulcus.

**Figure S2:**
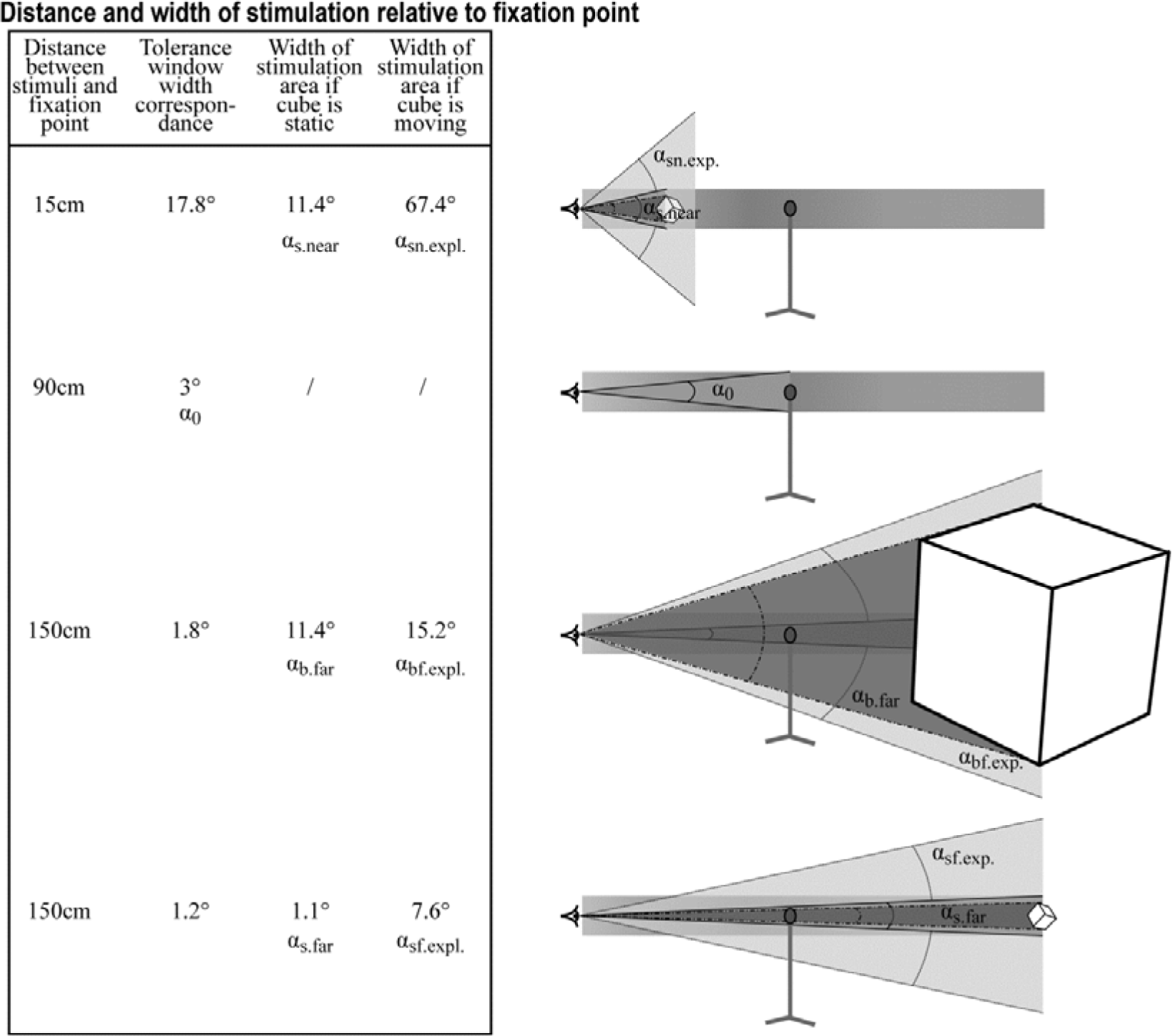
Graphical description of the expected range of eye movements for each type of stimuli if the monkeys had been fixating and tracking the objects rather than maintaining vergence onto the fixation spot. α_0_: angular range of eye movements when fixation is onα fixation LED; α_s.near_: angular range of eye movements if fixation were on static small near object; α_sn.expl_: angular range of eye movements if fixation were on dynamic small near object (i.e. matching the experimental design); α _b.far_: angular range of eye movements if fixation were on static big far object; α_bf.expl_: angular range of eye movements if fixation were on dynamic big far object (i.e. matching the experimental design); α_s.far_: angular range of eye movements if fixation were on static small far object; α_sf.expl_: angular range of eye movements if fixation were on dynamic small far object (i.e. matching the experimental design).

**Figure S3:**
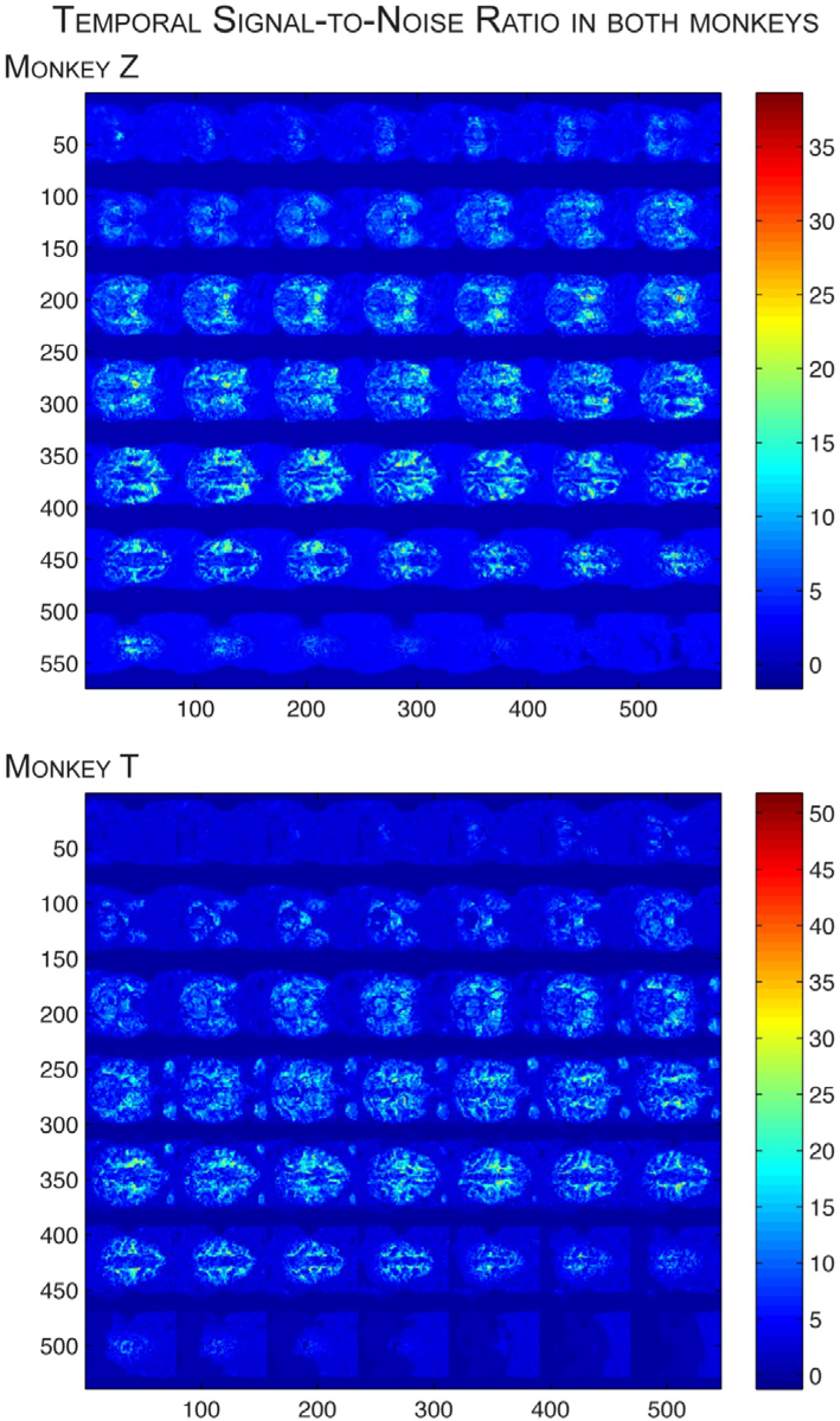
Temporal signal-to-noise ratio calculated in both monkeys through the brain, in transversal view.

